# Multiscale Analysis of *Candida albicans* Biofilms

**DOI:** 10.1101/2025.09.18.677201

**Authors:** Kai Li, Samantha Skivens, J. Edward F. Green, Alexander K. Y. Tam, Daniel R. Pentland, Hella Baumann, Campbell W. Gourlay, Benjamin J. Binder, Philippe P. Laissue

## Abstract

*Candida albicans* is an opportunistic fungal pathogen of significant biomedical concern. Its ability to colonize abiotic surfaces of clinical devices — such as catheters and airway management systems — can result in life-threatening sepsis, especially in immunocompromised patients. A deeper understanding of *C. albicans* biofilm development under different environmental conditions is essential for improving antifungal treatments. In this study, we examine *C. albicans* biofilm formation using live fluorescence microscopy across multiple scales and modalities, and introduce new quantification approaches. High-magnification tracking of hyphal tips reveals that hyphal elongation occurs intermittently rather than continuously. Using a new automated tracking approach, we show that hyphal emergence is initially rapid and slows down after approximately two hours. At lower magnifications, area coverage across large fields of view proves to be a robust and scalable metric. It is strongly influenced by seed density and extends analysis to later stages of growth. Elevated carbon dioxide levels significantly accelerate area coverage, promoting rapid biofilm expansion. Blue light illumination reduces *C. albicans* growth in a dose-dependent manner. Light-sheet imaging enables the long-term capture of vertical biofilm growth, complementing widefield-based approaches. We introduce logistic model parameters to effectively quantify the dynamics of surface area growth. The methodologies presented here are well-suited for high-content screening applications aimed at identifying compounds that inhibit or suppress fungal biofilm formation under clinically relevant conditions.

## 1 Introduction

Biofilms are microbial communities that adhere to surfaces and, over time, produce a protective extracellular matrix. While biofilms can play beneficial roles in processes such as wastewater treatment, they pose serious challenges in biomedical settings. A major challenge for example is the pathogenic yeast *Candida albicans*, which can colonize abiotic surfaces of airway management devices — including voice prostheses, tracheostomy tubes, and endotracheal tubing — leading to increased infection rates (Pentland et al. 2021). *Candida* responds to elevated levels of carbon dioxide (CO_2_), as found in the airways, by initiating biofilm formation. This in turn helps bacteria to become established on indwelling medical devices, increasing the risk of infections in patients (Pentland et al. 2020). By contrast, exposing *Candida* to blue light (BL) inhibits its growth and viability, potentially reducing biofilm formation and pathogenicity. Quantitative analysis is therefore crucial to better understand the fundamental mechanisms that inhibit or promote biofilm formation, and to develop improved models and methods for biofilm growth (Cámara et al. 2022).

Image-based analysis has become a powerful tool for studying biofilm development. Microscopy of fixed and labelled cells within biofilms has revealed complex internal structures, shedding light on the spatial organization and cell types present at specific time points. Live imaging extends this by capturing biofilm evolution over time, allowing researchers to track the location, orientation, shape, and progeny of cells and hyphae—from a single cell to a mature three-dimensional structure (Wong et al. 2021).

In bacterial systems, live imaging and analysis using dedicated open-source software (Hartmann et al. 2021; Heydorn et al. 2000; Vorregaard 2008) has revealed new growth modes (Qin et al. 2020). However, fungal biofilms require distinct imaging strategies due to their different growth patterns and architectures. For instance, Nagy et al. 2014 used brightfield microscopy to allow for high temporal resolution with minimal risk of causing phototoxicity. Kaneko et al. 2013 employed a side-view perspective to visualize developing *Candida albicans* biofilms. More complex and instrumentally demanding approaches using bespoke light-sheet setups have also been proposed by other groups in recent years (Gutiérrez–Medina et al. 2021; Licea-Rodriguez et al. 2019).

In this work, we introduce new approaches for the image-based analysis of *Candida albicans* biofilms, covering the full bioimaging pipeline — from sample preparation and image acquisition to image processing and visualisation. Using a multiscale approach, we examine the stages of biofilm formation, from biofilm initiation and the growth of single hyphae to immature and mature biofilms (Morelli et al. 2021). We employ different imaging modalities (conventional widefield fluorescence microscopy (WFM) and advanced light-sheet fluorescence microscopy (LSFM)) and magnifications ranging from 10*×* to *×* 90 . The new methods and metrics introduced here offer a comprehensive framework for quantifying and analysing biofilm formation, as summarised in the table below.

The methods and techniques introduced in this study demonstrate consistency across multiple scales of biofilm development and align with expected outcomes regarding the promotion and inhibition of biofilm formation by CO_2_ and BL, respectively. Our work provides an efficient pipeline for the visualisation and quantification of complex biofilm dynamics. This is an essential step toward the early diagnosis of fungal biofilm-related diseases and the development of effective strategies for their control and eradication.

## 2. Results

### Manual tracking of hyphal tips reveals that hyphal elongation is intermittent

To investigate the formation of *Candida albicans* biofilms during the earliest stages of development, we imaged individual yeast-form cells at 37^*°*^ C degrees using high magnification (Plan Apo *λ* 60*×* Oil, NA 1.42). A mitochondrial GFP tag (Duvenage et al. 2019b) was used to mark cell locations and to visualize hyphal emergence. Settled *Candida albicans* cells in their ovoid, unicellular yeast form served as the starting point for live imaging. Shortly after settling, these cells produced primary hyphae (Fig. 1A, arrowheads). The hyphae continued to grow from the yeast cells and could be reliably tracked for approximately five to seven hours. By the fifth hour, secondary hyphae branching from the still elongating primary hyphae became visible.

**Figure 1:**
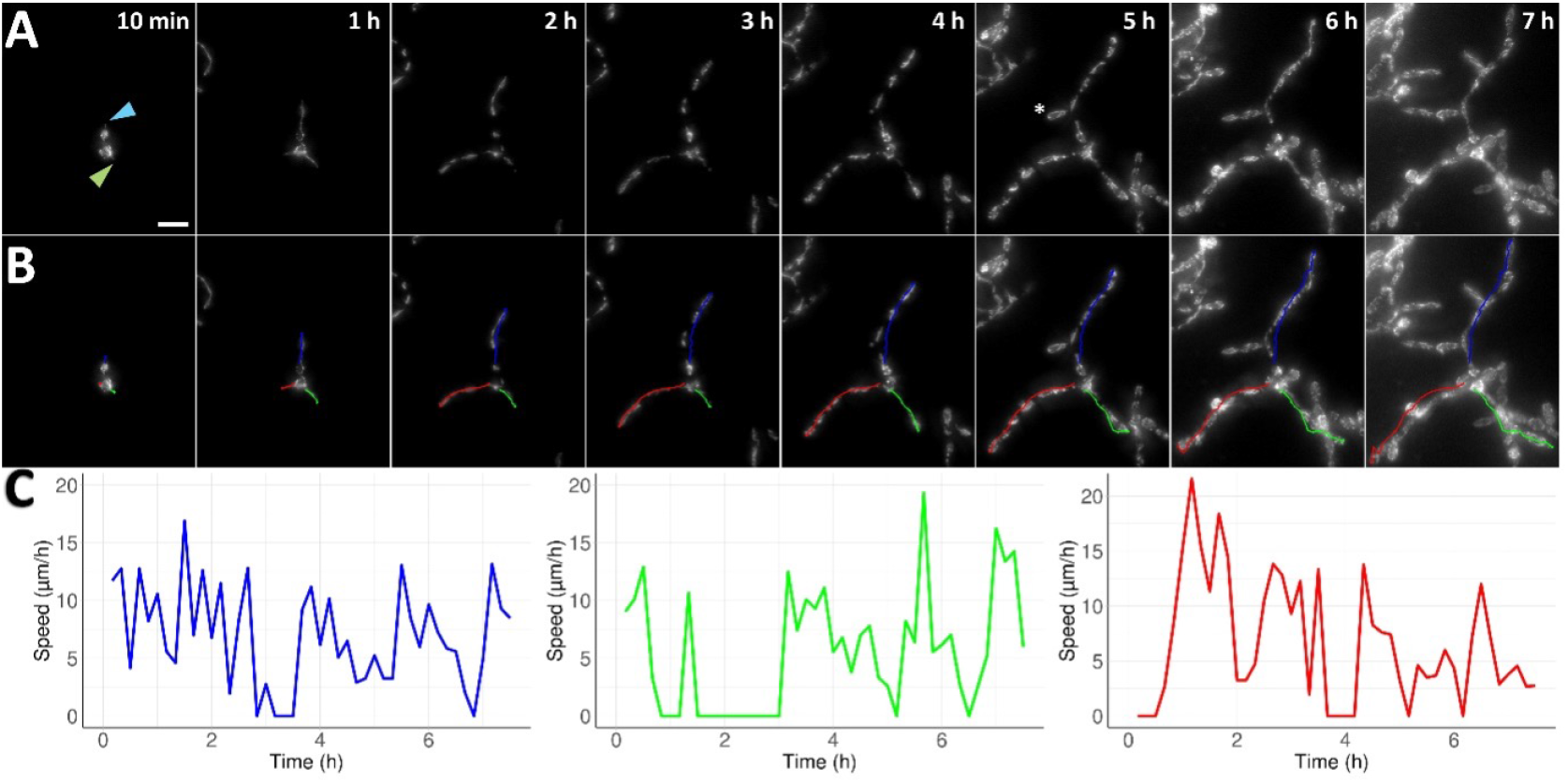
A) Greyscale images of *Candida albicans* in its initial unicellular yeast form, imaged at 60 *×* magnification. Hyphae emerge as early as 10 min after settling (light blue and light green arrowheads) and continue to elongate over the course of seven hours. At five hours (5 h), a secondary hypha (white asterisk) emerges from the primary hypha. Scalebar 10 µm. B) Paths of the emerging primary hyphae are shown (blue, green and red). C) Graphs of hyphal speed over seven hours, with colours corresponding to the hyphal tracks shown in B). Hyphal growth is intermittent, with phases of zero hyphal elongation.

We manually tracked hyphae by identifying the tip in sequential images (Schindelin et al. 2012). The individual trajectories of primary hyphae are visualized as coloured curves (blue, green, red) in Fig. 1B. The corresponding speeds are shown in Fig. 1C, revealing that hyphal elongation varies significantly over time and includes periods of stagnation, during which no elongation is observed.

This approach revealed that hyphal emergence is not a continuous process. In all tracked hyphae, we found that elongation is intermittent. This is not to say that no growth occurs during this time. Upon closer inspection, we observed thickening of the hyphal tip when hyphal elongation pauses. This suggests that growth continues during these periods, but in an isotropic way, rather than the polarised growth of hyphal elongation. This is, to the best of our knowledge, the first time that this intermittent hyphal extension, concomitant with isotropic thickening of the hyphal tip, has been described in *Candida albicans*.

By selecting trackable hyphae from twelve datasets with large fields of view, we extracted 23 hyphal trajectories with an average duration of 3.1 hours. The pooled speed measurements (Supplementary Fig. S1) reveal a general trend of hyphal elongation slowing over the course of five hours. However, manual tracking is time-consuming, susceptible to selection bias, and difficult to extend beyond the initial hours of hyphal development, as many hyphal tips become obscured in the dense network of neighbouring hyphae. To address these limitations, we developed an automated method for hyphal tracking.

### Automated hyphal tracking reveals that hyphae emerge quickly, then slow down

We developed a novel algorithm to track hyphal tips, enabling efficient, large-scale, and unbiased tracking over extended periods (see Methods). This “big-data” approach allows for the rapid analysis of hundreds of hyphal trajectories without observer-related bias. While manual selection and tracking required approximately two working days to generate 23 trajectories with a total length of 979 µm, automated tracking produced 181 trajectories spanning 3983 µm within minutes. Segmenting the first 7.5 hours of hyphal development into three consecutive 2.5-hour time windows, we plot the displacement of the tracked paths (*i*.*e*., distance) versus time *t* in Fig. 2A–C. For each time window, the corresponding average displacement data are plotted as solid curves in Fig. 2D. Shaded regions represent the 95% confidence intervals, calculated as 1.96 times the standard error.

**Figure 2:**
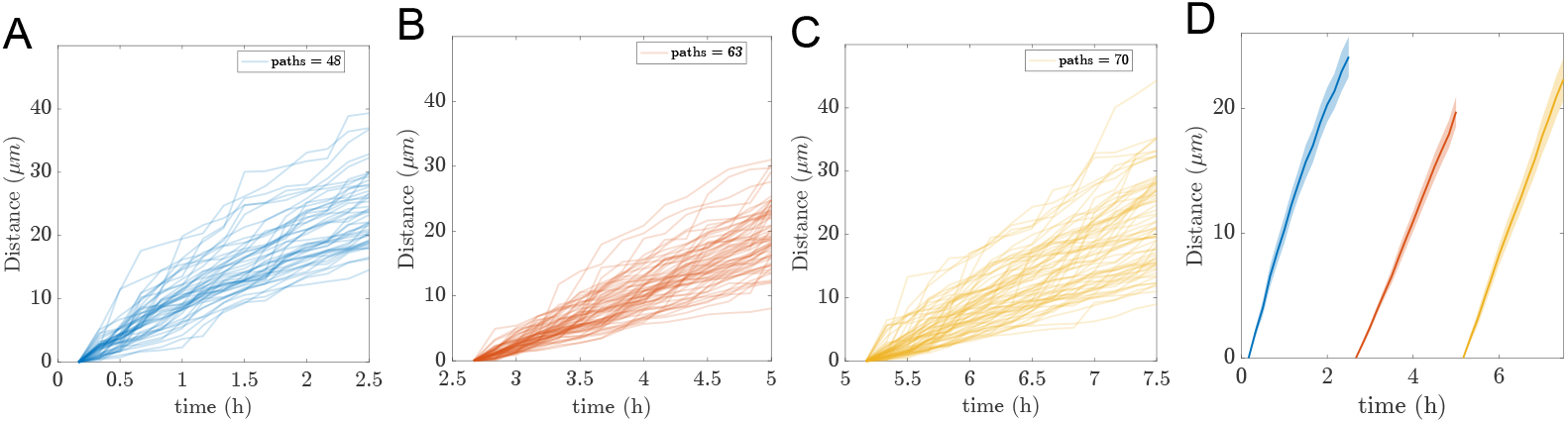
Automated tracking of multiple hyphae in three time windows of 2.5 h length each. A) to C) Hyphal path distance with time. A) 0–2.5 h, 48 hyphae. B) 2.5–5 h, 63 hyphae. C) 5–7.5 h, 70 hyphae. D) The time windows from A to C are concatenated in one graph and shown as average distance (µm) over time. The semi-transparent region around each solid curve for the mean represents the 95% CI.

The average displacement data exhibit a linear trend within each of the three 2.5-hour time windows shown in Fig. 2D. Accordingly, we fit a linear regression model of the form

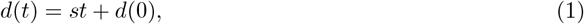

to the average data in each window using the `polyfit()` function in MATLAB. The slope parameter *s* provides an estimate of the mean speed during each time interval. Each fit yielded a near-perfect linear relationship, with *R*^2^ = 0.99 across all three time windows, which aligns with the tight CI shown in Fig. 2D.

For each consecutive 2.5-hour time window, we obtain the following estimates for the average speed, *s* = 10.34, 8.47, 9.59. Given the near-perfect linear fits and narrow confidence intervals, we infer that hyphae initially emerge at high speed, slow down during the middle phase, and then accelerate again. Through this computational, data-driven analysis, we have uncovered a previously undescribed pattern in hyphal emergence.

### Fluorescent mitochondria indicate cellular stress at high magnification

High magnification is required to enable the tracking of single hyphae. But higher magnification also increases the light intensity (strictly speaking the irradiance) that the sample is subjected to. It should be noted that a two-fold increase in magnification leads to a four-fold increase in irradiance (if using the same illumination power for each magnification). We thus monitored live imaging at high magnification to check for phototoxic effects. Image acquisition at magnifications higher than 60*×* induced mitochondrial fragmentation (Fig. 3 A,B). Mitochondrial fragmentation is a known indicator of phototoxicity (Higuchi-Sanabria et al. 2016; Kiepas et al. 2020). However, mitochondrial width remained unchanged even at 150 *×* magnification (Fig. 3 C), indicating that the observed fragmentation was not a result of image processing artifacts such as loss of fluorescent intensity through photobleaching. These observations support the use of our *C. albicans* strain with fluorescently labelled mitochondria as a reliable tool for monitoring cell health. They also suggest that hyphal tracking should be restricted to the early stages of biofilm development to minimise phototoxic effects.

**Figure 3:**
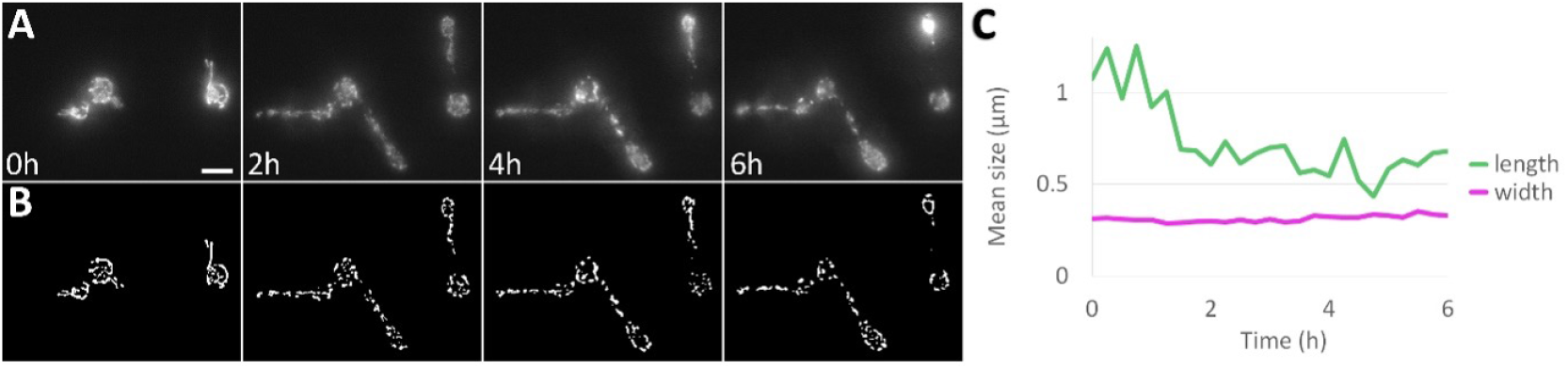
Mitochondrial fragmentation indicates phototoxicity at high magnification. A) Fluorescently labelled mitochondria show hyphal emergence from single cells at 150 *×* magnification over six hours. Scalebar 5 µm. B) Processed images showing the isolated mitochondria. C) Quantification showing mitochondrial fragmentation. The average length (green) of mitochondria is greatly reduced after two hours, while their width (magenta) remains unchanged.

We have demonstrated the utility of high-magnification imaging for studying the early stages of *C. albicans* biofilm formation, enabling visualization and quantification of cellular settlement and hyphal elongation. Our results show that hyphal elongation is intermittent, and we introduce a novel method for automated measurement of elongation dynamics, revealing a decline in mean hyphal elongation speed over time. However, this high-resolution approach is inherently limited: it cannot be reliably extended beyond the first 5 to 7.5 hours of development and requires careful control to avoid phototoxic effects that may compromise data quality and interpretation. To overcome these limitations and enable more robust, long-term quantification of biofilm growth, we pursued an alternative imaging strategy.

### Progressive surface area coverage as a metric for mature biofilms

While measuring the speed of hyphal emergence provides valuable insights into early-stage biofilm development at high resolution, it is not a robust metric for quantifying growth in mature biofilms. Tracking becomes unreliable when individual hyphal tips are no longer clearly distinguishable, and as biofilms mature, they increase in density and accumulate extracellular matrix. Although hyphae continue to grow through this matrix, their elongation speeds are influenced by both the matrix and the proximity of neighbouring hyphae. This is shown in Fig. 4(A-C). The square yellow insets are magnified in Fig. 4(D-F).

**Figure 4:**
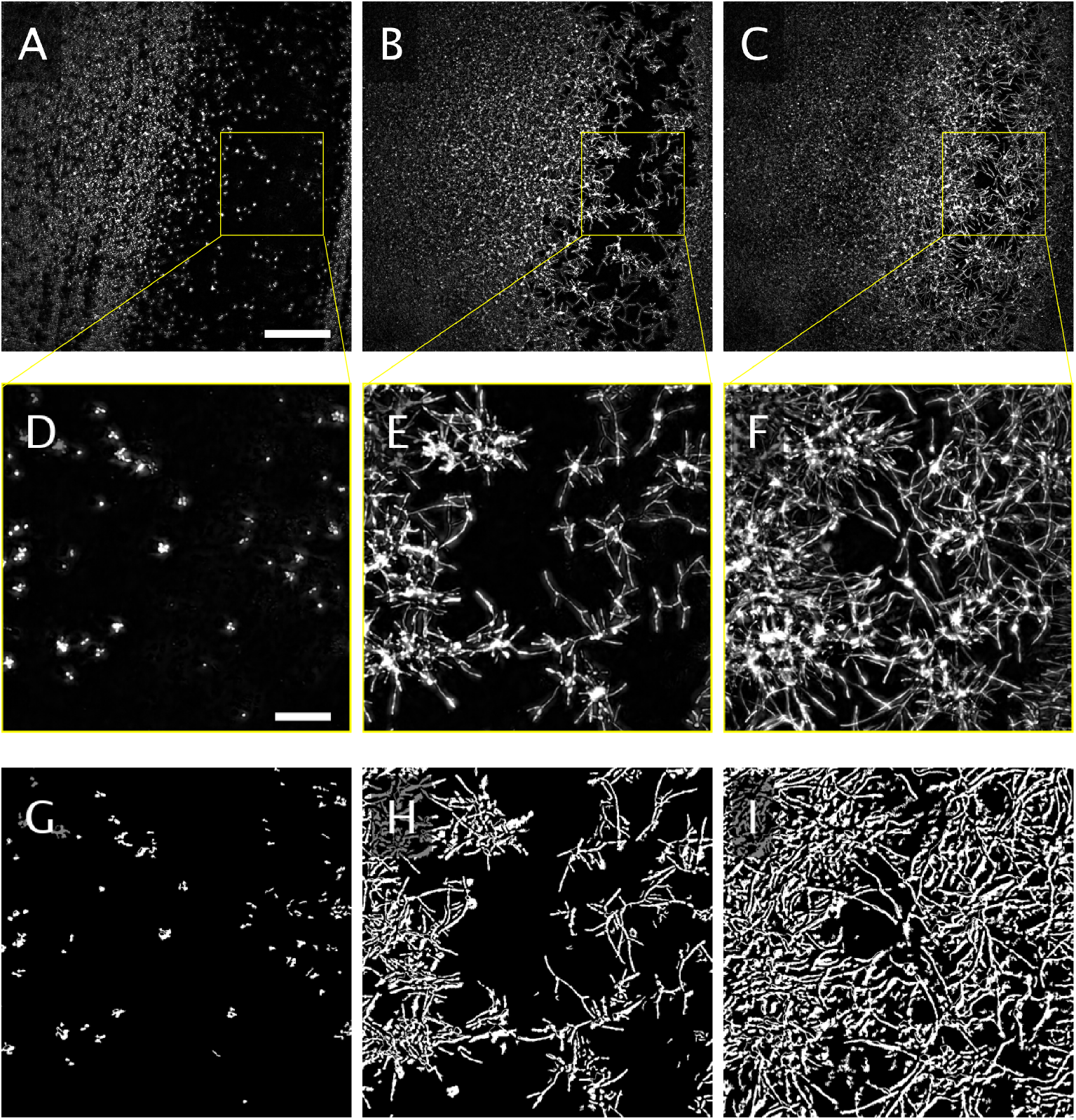
Large FOV at low resolution showing A) biofilm initiation at single cell stage with scalebar 200 µm, B) biofilm development after 6 hours and C) after 12 hours. D) to F) are the magnified insets of the yellow squares in A) to C), respectively, with a scalebar of 50 µm. G) to I) are the corresponding binary images.

A more suitable metric for long-term growth analysis is the progressive coverage of surface on which the biofilm is growing. This approach enables tracking biofilm expansion from individual cells to mature, densely packed communities. Because it does not require the resolution necessary to follow individual hyphae, lower magnifications can be used. This further minimises the risk of phototoxicity and also provides a larger field of view (FOV). For example, the FOV at 60 *×* magnification (using a large camera ROI of 2394*×* 2394 pixels) spans 259.4 µm per side, covering an area of 0.067 mm^2^. By comparison, a 15*×* FOV (using the same ROI) spans 1063.1 µm per side, covering 1.13 mm^2^, an area nearly 17 *×* larger. This is important for the analysis of local differences in biofilm growth. However, in large FOVs at low magnification, fluorescence intensity decreases towards the edges and corners, a common microscopy artefact known as ‘vignetting’. For this reason, adaptive thresholding must be used to compensate for the lower signal-to-background ratio at image edges and corners (Fig. 4(G-I)).

### A scratch assay enables variable seed densities at the start of a time-lapse recording

We developed a scratch assay for *C. albicans* biofilms to examine how the number of single yeast cells at the start of an experiment (*i*.*e*. the seed density) affects the rate of surface coverage over time. Typical initial conditions are shown in Fig. 5A, which displays a large FOV imaged at 15 *×* magnification. Differences in seed density across the image result from the scratch assay. To quantify these differences, we partitioned the image into a 8*×* 8 grid and classified each square’s initial density as either sparse (red markers) or dense (blue markers), using 5% surface coverage as the threshold between the two categories. After 12 hours, the biofilm covers most of the surface within the FOV (Fig. 5C). We further show that the number of cells at the start of a time-lapse recording can be accurately determined, as the measured area is directly proportional to the total cell count (Supplementary Fig. S2).

**Figure 5:**
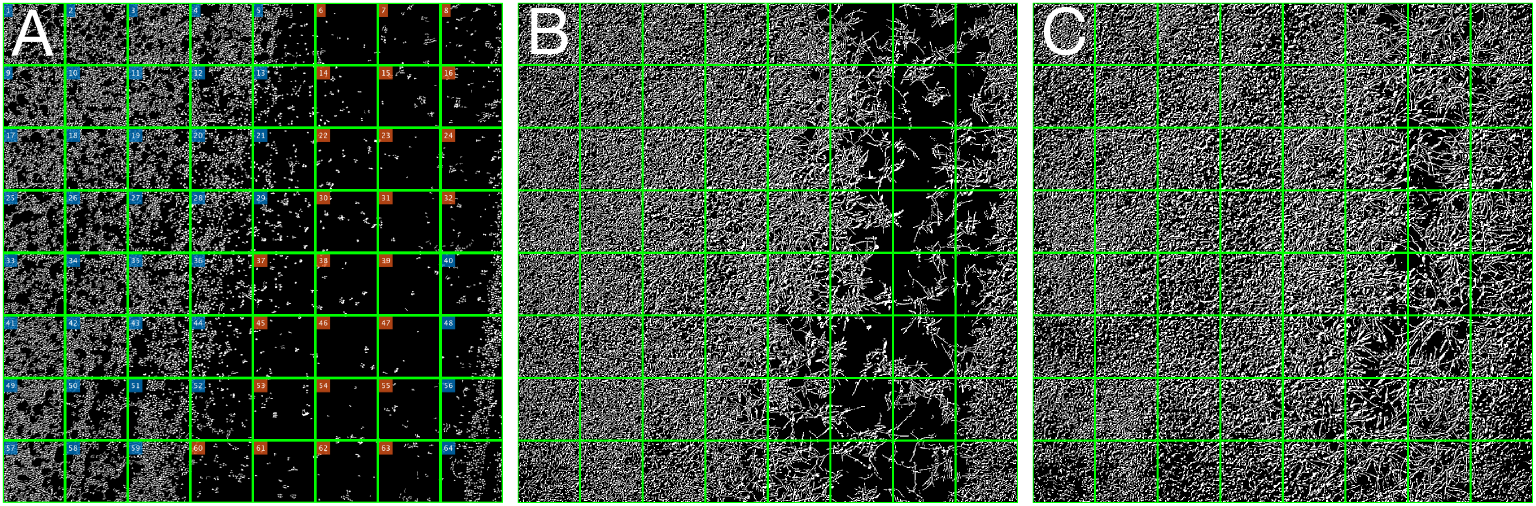
Biofilm scratch assay. A) Initial condition. The FOV imaged at 15 *×* magnification partitioned into 64 squares. High initial density, blue markers. Sparse initial density, red markers. Coverage of basal well surface after B) 6 hours and C) 12 hours. Each partitioned square covers 132 µm *×* 132 µm.

### Surface coverage rate is influenced by the seed density

Surface coverage resulting from biofilm growth is quantified i Fig. 6A. The progression of surface coverage for each individual grid partition (as defined in Fig. 5) are shown as individual curves, categorised by the two seed densities (sparse (yellow) and dense (cyan)). Using the same colour coding, i Fig. 6B shows their mean value as solid curve with 95% CI (semi-transparent shading). This shows that the rate of surface coverage is influenced by the seed density. Seed density must thus be taken into account when determining growth rates. It should also be similar when comparing different experimental conditions.

**Figure 6:**
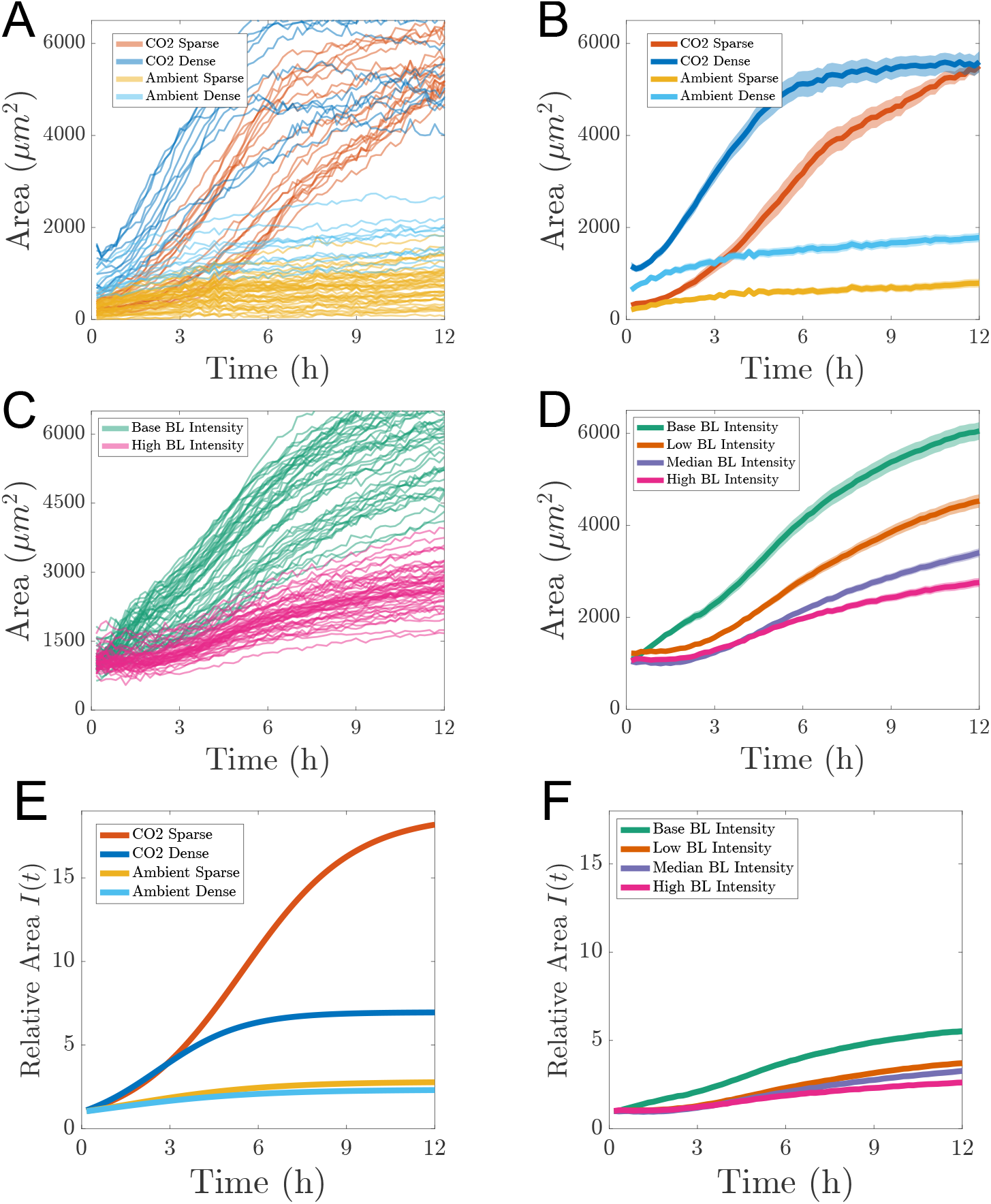
Quantification of progressive surface coverage for different experimental conditions and factors. A) Different CO_2_ conditions combined with varying seed densities. Elevated CO_2_ and dense seed density (dark blue). Elevated CO_2_ and sparse seed density (red). Ambient CO_2_ and dense seed density (cyan.) Ambient CO_2_ and sparse seed density (yellow). Individual curves represent measurements for image partitions in Fig. 5. B) Combinations and colour-coding as in A. Mean of all partitions (opaque curves) with 95% CI (semi-transparent shading). C) Comparison of biofilm growth at different BL intensities. Base BL intensity (green curves). High BL intensity (magenta). The different curves represent individual measurements from a partitioned FOV for each intensity treatment. D) Combinations and colour-coding as in C, with additional BL intensities. Low BL intensity (orange). Medium BL intensity (purple). Mean (opaque curves) with 95% CI (semi-transparent shading). E) and F) Curve fitted relative area coverage, combinations and colour-coding as in B and D. KL: Update relative area to *S*(*t*).

### Elevated CO_2_ levels increase the surface coverage rate

Physiological levels of CO_2_ (as opposed to the lower ambient levels) have been shown to accelerate the growth of *C. albicans* (Pentland et al. 2021). We imaged C. albicans surface coverage and quantified it, additionally taking into account the aforementioned seed density. This is shown in Fig. 6A and B. Under ambient CO_2_ conditions, the high seed density leads to a biofilm surface coverage approximately twice that of the sparse seeding condition (cyan and yellow curves in Fig. 6(A,B). Notably, neither condition results in the surface being fully covered after 12 hours. A similar trend is seen under elevated CO_2_ levels for both sparse and high initial densities (red and blue curves, <5 h). However, the rate of coverage is substantially higher under elevated CO_2_. Specifically, the dense initial density condition reaches confluence at approximately 5 hours, while the sparse condition does so by around 12 hours. These observations indicate that elevated CO_2_ levels have a more pronounced effect on biofilm growth than initial seeding density, promoting significantly more rapid biofilm development.

In addition to seeding density and CO_2_ levels, other factors such as temperature and BL exposure can influence the rate of area coverage. For example, environmental conditions that mimic those found on medically relevant abiotic surfaces—such as airway management tubes—include a temperature of 37^*°*^C (body temperature) and elevated CO_2_ levels (5 % above ambient). In the following experiments, we replicate these clinically relevant growth conditions and apply increasing BL intensities to examine the effect on surface coverage rates.

### Increasing BL intensity reduces the surface coverage rate in a dose-dependent manner

Using similar cell densities as starting condition, we increased BL illumination intensities from a non-invasive baseline to low, medium and high levels. Exposure times were kept constant. The resulting analysis is presented in Fig. 6(C,D). BL shows a dose-dependent inhibitory effect on *C. albicans* biofilm growth. These results corroborate surface coverage rate as a robust metric to quantify factors promoting as well as reducing *C. albicans* biofilm growth. The additional advantage is that large FOVs are assessed using WFM, a widespread, gentle and highly affordable fluorescence microscopy modality, in combination with simple and rapid image processing steps.

### Estimate of growth rate and long-term limit of surface area coverage

We observe that the trend in the average area coverage data (Fig. 6B and D) follows a logistic pattern and can be modeled by

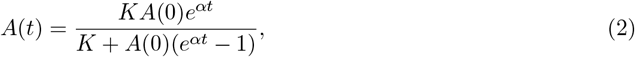

where *α* is the growth rate, *K* represents the long-term limit of the coverage area (which we will call the carrying capacity), and *A*(0) is the initial area coverage or seed density. The logistic curve (2) was fitted to the average area data, using the `lsqcurvefit` function in MATLAB, yielding estimates for the parameters *α, A*(0) and *K*. The coefficient of determination values, *R*^2^ *≈* 1, suggests a near perfect fit across all experiments (see Table 2).

**Table 1:**
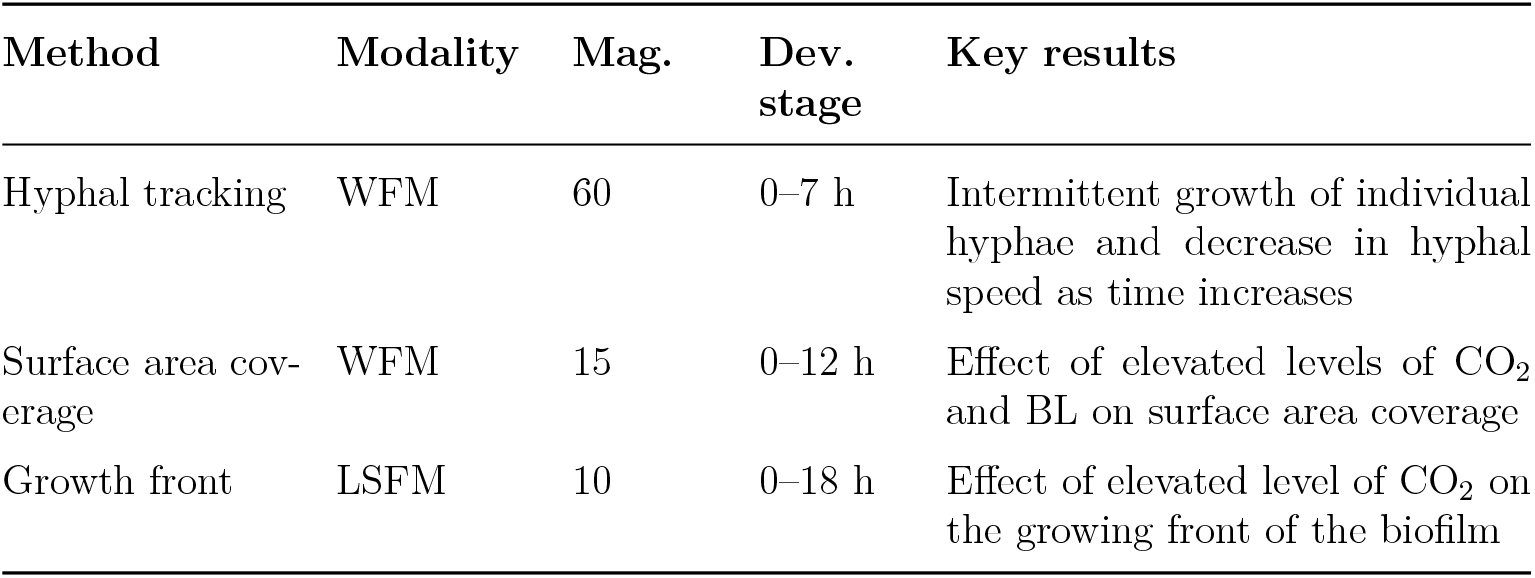
Summary of imaging methods and analysis for biofilm quantification. *Mag*. Magnification, *Dev*. Developmental.

**Table 2:** Values of the parameters for the logistic curve best-fit as computed in MATLAB using lsqcurvefit().

| | $\alpha$ | $A(0)$ | $K$ | $\alpha/K$ | $K/A(0)$ | $R^2$ |
| --- | --- | --- | --- | --- | --- | --- |
| CO<sub>2</sub> Sparse | 0.53 | 290 | 5447 | $9.73 \times 10^{-5}$ | 18.78 | 0.99 |
| CO<sub>2</sub> Dense | 0.69 | 798 | 5555 | $1.24 \times 10^{-4}$ | 6.96 | 0.99 |
| Ambient Sparse | 0.42 | 261 | 733 | $1.61 \times 10^{-3}$ | 2.81 | 0.95 |
| Ambient Dense | 0.40 | 744 | 1735 | $5.72 \times 10^{-4}$ | 2.33 | 0.98 |
| Base BL Intensity | 0.36 | 1042 | 6434 | $5.60 \times 10^{-5}$ | 6.17 | 0.99 |
| Low BL Intensity | 0.27 | 908 | 5455 | $4.95 \times 10^{-5}$ | 6.01 | 0.99 |
| Medium BL Intensity | 0.26 | 749 | 4092 | $6.35 \times 10^{-5}$ | 5.46 | 0.99 |
| High BL Intensity | 0.23 | 870 | 3256 | $7.06 \times 10^{-5}$ | 3.74 | 0.99 |

Additionally, we define the relative area coverage as

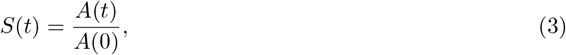

for each experiment. Scaling the area *A*(*t*) by the initial area *A*(0) allows us to examine biofilm growth (*i*.*e*. surface coverage) relative to its starting value, providing a metric that enables simultaneous visualisation of relative growth across all experimental conditions. One striking feature of this comparison is the amplifying effect of CO_2_ in conditions of sparse seed density, as evidenced by the red curve in Fig. 6(E).

The estimated parameters *α, A*(0), and *K* characterise and quantify the growth dynamics across all experiments (see Table 2). As expected, we observe higher values of the intrinsic growth rate *α* in the CO_2_-enriched experiments compared to those conducted under ambient conditions. Similarly, and in line with expectations, *α* decreases with increasing BL intensity. To quantify long-term growth, we evaluate the scaled area at saturation (i.e. the long-term limit of surface coverage) using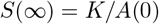, which represents the fold increase in area relative to the initial surface coverage. For instance, under CO_2_-enriched conditions with sparse initial density, the surface coverage increased approximately 18-fold. The ratio *α/K* quantifies how rapidly the area approaches its carrying capacity. Among the experimental conditions, the Ambient Sparse setup reaches its maximum area most quickly, whereas the CO_2_ Sparse and High BL intensity conditions exhibit the slowest progression toward their carrying capacities. This analysis demonstrates the utility of the logistic model parameters in effectively quantifying and describing the surface area growth dynamics.

### LSFM enables long-term imaging of biofilm growth, including its vertical component

The methods and metrics discussed thus far have focused on conventional, single-objective WFM using an inverted microscope. This modality is advantageous due to its widespread availability in laboratories and imaging facilities focused on biomedical research. However, a key limitation of inverted WFM is that once the imaged surface (in our case the basal well surface) becomes fully covered by hyphae, further biofilm development cannot be visualized.

To address this limitation, we imaged biofilm growth from above to capture upward movements of hyphae (*i*.*e*. hyphae extending at an angle, moving away from the basal surface of the well). Mature fungal biofilms are known to exhibit substantial vertical growth (Morelli et al. 2021), which cannot be captured with inverted WFM. Using LSFM in an upright, dual-objective configuration (*i*.*e*. diSPIM (Kumar et al. 2014; Wu et al. 2013)), we were able to visualize and quantify biofilm development in this axial direction. This imaging modality requires a distinct processing pipeline, as outlined in Fig. 7.

**Figure 7:**
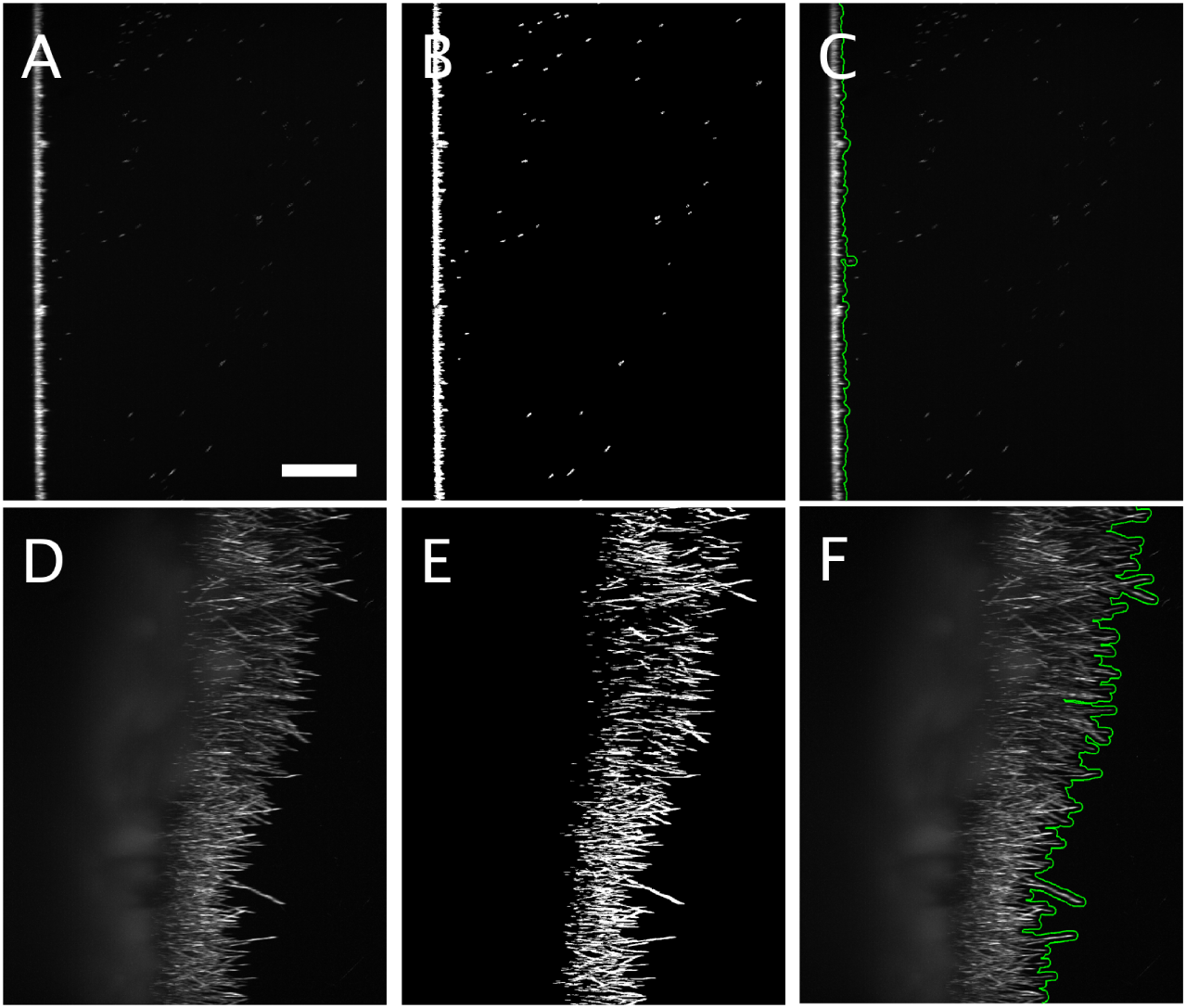
Vertical expansion of a *C. albicans* biofilm using LSFM. The top row shows images at the time point 0, the bottom row shows images after 18 h of growth. (A,D) Original images. (B,E) Binarised images after thresholding. (C,F) Processed images showing the biofilm front (green curves) of growing hyphae. Scalebar 200 µm.

LSFM enables detailed imaging of biofilm structures in three spatial dimensions over time. As an inherently gentle imaging modality, LSFM significantly reduces phototoxicity compared to conventional fluorescence techniques with optical sectioning. However, this advantage comes with the trade-off of generating large volumes of data. To make image processing more tractable, we collapsed the third spatial dimension by computing a maximum intensity projection.

Using this projection, we identified the leading front of the growing hyphae and quantified its progression by subsampling rectangular regions or partitions of interest. By pooling two replicates for each condition (ambient and elevated CO_2_, both at 37°C), we observed that elevated CO_2_ conditions promote faster biofilm growth (as expected), as indicated by greater front-line displacement over the same time interval (Fig. 8).

**Figure 8:**
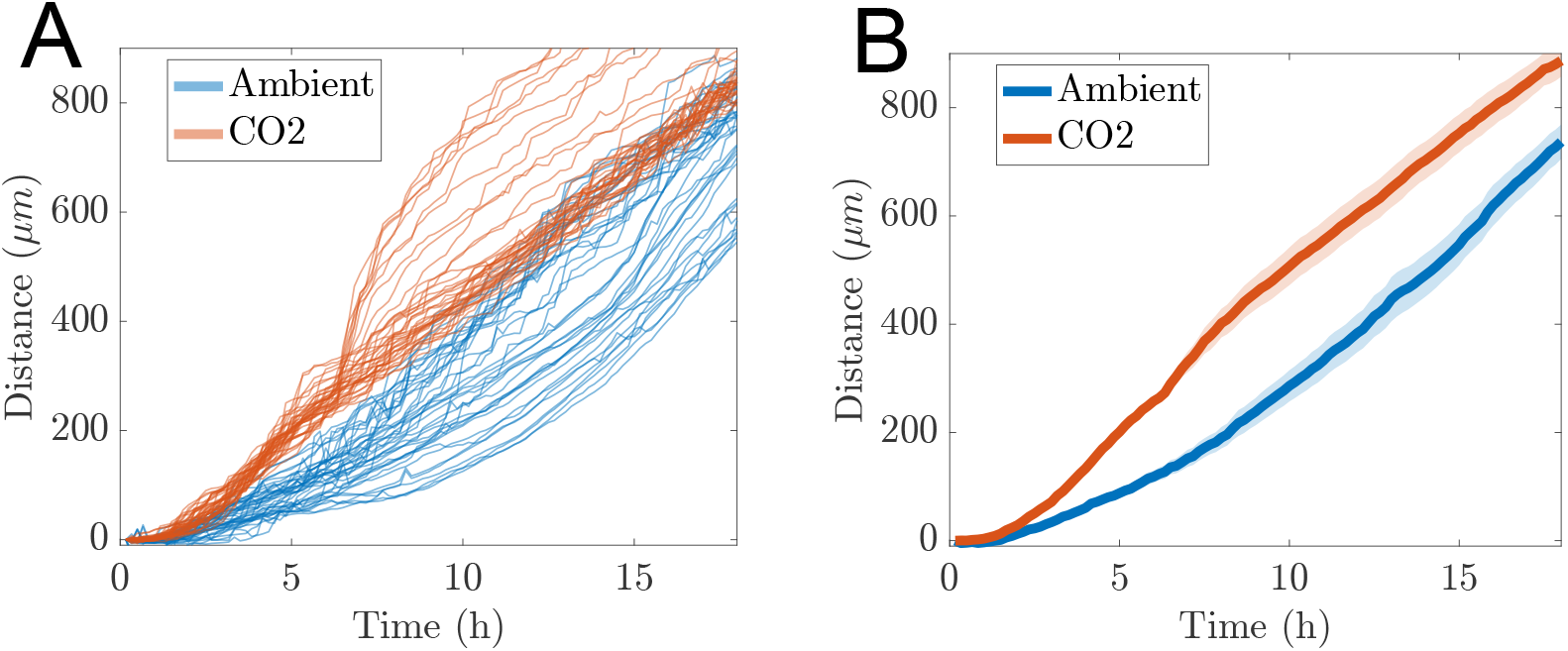
Time evolution of mature biofilm front in direction perpendicular to basal well surface. Ambient conditions, blue. Elevated CO_2_ levels, red. A) Displacement of individual equally partitioned regions of the front. B) Average displacement of front (solid curves) with the semi-transparent region indicating the 95% CI.

Unlike the inverted WFM approach used previously, LSFM allows imaging from above, enabling continuous observation of biofilm development even after complete coverage of the substrate. As a result, the growth curves in Fig. 8 do not plateau but instead show ongoing vertical expansion of the biofilm.

### Estimating the speed of the growing biofilm front

Similar to the displacement data from hyphal tracking, we observe a linear trend in the time evolution of the average displacement of the biofilm front (see Fig. 8(B)). Therefore, we fit the linear model given in Eq. (1) and obtain speed estimates of *s* = 54.23 µm h^*−*1^ (*R*^2^ = 0.97) for the CO_2_-enriched experiment and *s* = 43.10 µm h^*−*1^ (*R*^2^ = 0.99) under ambient conditions. Thus the average speed of the LSFM front is another metric to quantify different growth dynamics. Note that rough estimates of the speed can be obtained by simply dividing the displacement at the final time point by the total duration of the experiment, yielding *s* = 49.27 µm h^*−*1^ and *s* = 40.94 µm h^*−*1^ for the CO_2_-enriched and ambient conditions, respectively.

## 3 Discussion

### Hyphal tracking at high magnification for early stages of biofilm development

Tracking hyphal tips at high spatial resolution provides valuable insights into hyphal elongation dynamics. We show that hyphal extension is intermittent, alternating between phases of active growth and pauses. These pauses frequently coincide with the formation of bulbous tips, a phenotype observed in various filamentous fungi when polarized growth is disrupted (Riquelme et al. 2018). Whether this alternating growth behaviour represents a programmed mechanism for reassessing growth direction (e.g., in response to nutrient gradients) remains an open question.

Manual tracking of hyphae is labour-intensive and introduces a bias toward selecting clearly identifiable tips in less crowded regions. In contrast, our development of an automated tracking algorithm enabled the analysis of a large number of elongating hyphae in a short time, free from user selection bias. This revealed a longer-term pattern of hyphae slowing down over time. High-throughput tracking is especially advantageous in the context of high variance due to the intermittent nature of hyphal elongation.

However, hyphal tracking is limited to the early stages of biofilm development and becomes unreliable beyond 5 to 7.5 hours after hyphal emergence. This is consistent with previous work by Nagy et al. (2014), who restricted their analysis to the first five hours. Our measured hyphal extension speeds are broadly in agreement with values reported in prior studies (Chen et al. 2020; Kaneko et al. 2013; Nagy et al. 2014; Sevilla et al. 1986). Nonetheless, comparisons are limited by differences in *C. albicans* strains, culture media, and incomplete methodological reporting in many of those studies. In particular, seed density is rarely accounted for. While hyphal elongation speeds provide valuable mechanistic insights at the single-cell level, the inherent variability of intermittent growth makes averaging across hyphae a limited metric for assessing overall biofilm development.

### Coverage of the basal well surface provides robust and accessible metrics

Area coverage measurements offer a straightforward alternative to single-hypha tracking and can be conducted at lower magnification, capturing a much larger sampling area. This allows extension of the observation period up to the point of full FOV coverage. In the case of single-species *C. albicans* biofilms, this extends analysis windows from 7.5 hours (tracking) to up to 12 hours (coverage). The slope of the area coverage curve reflects the rate at which the biofilm spreads over the substrate and serves as an accessible and informative metric.

We find that seed density is a critical parameter for both area coverage and biofilm growth more generally. The influence of inoculum size has been described previously for *Ceratocystis ulmi*, attributed to quorum sensing effects (Hornby et al. 2004). In *C. albicans*, quorum sensing is mediated by farnesol (Hornby et al. 2001; Ramage et al. 2002). To address local variation in seed density within the FOV, we employed image subsampling (Fig. 5), which also provides additional datapoints for statistical analysis. In cases of high seed density where growth saturates rapidly, we implemented a scratch assay (see Methods) to locally reduce cell numbers, allowing more meaningful observation and quantification.

One limitation of area coverage analysis using an inverted microscope is that growth beyond full FOV coverage cannot be assessed. In contrast, imaging from above with an upright microscope allows continued observation of vertical hyphal growth at later stages. We validated the area coverage method by demonstrating a dose-dependent inhibitory effect of blue light on *C. albicans* growth. In this context, phototoxicity is intentional and could potentially be enhanced through the use of photosensitizers. Given the clinical relevance of *C. albicans* biofilms on abiotic surfaces such as airway management devices and catheters, such approaches have significant translational potential.

### Advantages of using conventional WFM

Live fluorescence microscopy enables highly selective labeling of proteins and structures of interest. The strong inherent contrast between signal and background facilitates efficient image segmentation compared to other optical contrast techniques such as differential interference contrast or phase contrast. By acquiring image stacks in three spatial dimensions over time, we capture biofilm structures both on and above the substrate. For analysis, we collapse the *z*-axis using maximum intensity projections, simplifying processing and reducing data volume while retaining meaningful spatial information.

For detailed structural imaging with optical sectioning, confocal modalities such as spinning disk or laser-scanning microscopy are available. However, due to their higher phototoxicity, these techniques are better suited to endpoint analyses (Morelli et al. 2021; Pentland et al. 2020).

WFM remains one of the most accessible fluorescence imaging techniques and is well-suited for live imaging due to its gentle illumination. This advantage is further enhanced by modern scientific cameras (*e*.*g*. sCMOS), which combine high quantum efficiency in the visible range (400 nm to 800 nm) with low read noise, enabling effective imaging under low-light conditions. Nonetheless, phototoxicity can still occur, especially at high resolutions. We show that it manifests rapidly, emphasizing the need to keep imaging resolution as low as possible while still capturing the relevant biological information.

This supports the use of area coverage as a robust metric, as it enables low-magnification imaging with large FOVs. The combination of area coverage analysis, large FOV, low magnification, image subsampling, and multiwell plate imaging on an automated XY stage facilitates high-content screening. This approach is well-suited for both industrial and biomedical applications, including drug discovery and toxicity testing aimed at identifying compounds that inhibit or suppress biofilm formation under diverse environmental conditions.

LSFM is a more demanding technique in terms of sample preparation, image acquisition and image processing. However, its upright configuration and inherently low phototoxicity enable the long-term capture of vertical biofilm growth, which is a crucial component in mature biofilms. LSFM is thus an indispensible modality to complement the imaging of WFM-based biofilm development.

## 4. Conclusion

We present an integrated set of methods for quantifying *Candida albicans* biofilm growth, encompassing sample preparation, image acquisition, image processing, and quantitative analysis. Our approach offers flexible and complementary metrics that are tailored to different spatial resolutions and stages of biofilm development. For early-stage growth, high-magnification imaging enables analysis of hyphal extension dynamics, while at later stages, area coverage at low magnification provides a robust, scalable measure of biofilm expansion.

Among the metrics evaluated, area coverage proved to be a particularly effective proxy for biofilm growth, with clear dependence on inoculum density, CO_2_ conditions and BL intensity. The coverage of the basal well surface offers a convenient and interpretable point of comparison across experiments. At higher magnifications, we also note increased background fluorescence and other photophysical effects, underscoring the need to balance resolution with imaging constraints such as phototoxicity.

Finally, our findings raise an intriguing possibility for future work: Can hyphal growth dynamics be inferred from area coverage measurements alone? If so, this would offer a powerful tool for large-scale, high-throughput analysis of fungal biofilms.

## 5 Methods

### Sample preparation

*C. albicans* (strain SN250, TFKO Library Reference Strain (Homann et al. 2009)) transfected with a mitochondrial-target GFP (Duvenage et al. 2019a) and maintained on an agar plate, was used in this study. For each experiment, a stock solution was made by scraping a small amount of *C. albicans* from the agar plate using a clean pipette tip, and adding it to a sterile test tube with 10 ml Lonza RPMI 1640 growth medium. The test tube was shaken on a vortex mixer to suspend the cells. 500 µl stock solution was added to each well of an ibidi 8-well high ibiTreat plate.

### Scratch assay

To create FOVs with varying seed density, a scratch assay was construed. This involved using a clean pipette tip to scratch the bottom of each well in the 8-well plate in all directions. 500 µl of resuspended cells in growth medium was then removed from each well, and a clean pipette tip used to add 500 µl fresh growth medium back into each well.

### Fluorescence microscopy and imaging conditons

Widefield fluorescence microscopy was performed using a Nikon ECLIPSE Ti2-E Inverted Microscope with a motorized XY and Z stage and a 25 mm field of view with a 1.5*×* tube lens (Nikon Corporation, Tokyo, Japan). The microscope was equipped with a Crest X-Light V3 Spinning Disk Confocal (Crestoptics S.p.A., Rome, Italy), but used in widefield fluorescence mode. The light source used was an LDI-7 Laser Diode Illuminator (89 North) with TTL triggering in a Ubob42 NIDAQ Ultimate Breakout Box to avoid illumination overhead. Images were acquired at 16-bit using a Teledyne Photometrics Kinetix camera (Tucson, AZ, USA). Image acquisition was controlled with NIS-Elements AR (v5.42.02, Build 1801) on an HP Z4 High-End workstation with Intel Xeon 3.9GHz and NVIDIA Quadro RTX4000. GFP was excited at 470 nm using hard-coated interference filters (Semrock Inc., IDEX Corp., IL, USA). Objectives used were Nikon CFI PlanFluor 10 *×* (NA 0.3, WD 16 mm), CFI Plan Apochromat Lambda D 20X (NA 0.8, WD 0.8 mm) and CFI Plan Apochromat Lambda D 60X Oil (NA 1.42, WD 0.15 mm), often in combination with the 1.5x tube lens. Inoculated ibidi 8-well microplates were placed inside an environmentally controlled Okolab cage incubator on the microscope stage. Samples were incubated at 37 ^*°*^C, with manual gas mixing set to low airflow (0.6 litres/minute) and using CO_2_ concentrations of 0.5 (‘ambient’) and 5% (‘CO_2_’). Each field of view (2048 *×* 2048 pixels) was captured as a shallow z-stack (with 5–9 focal planes) and collapsed into a maximum intensity projection to include, where applicable, vertically growing components of a biofilm. The Perfect Focus System (v4) ensured a stable focus between images, and Nikon JOBS (2.0) enabled variable illumination between wells. For BL exposure experiments, 100 µmol m^*−*2^ s^*−*1^ was used as baseline intensity, followed by 330 µmol m^*−*2^ s^*−*1^ (‘low intensity’), 1000 µmol m^*−*2^ s^*−*1^ (‘medium intensity’) and 3000 µmol m^*−*2^ s^*−*1^ (‘high intensity’), using 900 milliseconds exposure time and 15x magnification in each case.

Light-sheet microscopy was conducted on an Intelligent Imaging Innovations (3i) Marianas LightSheet system (MLS) inside an environmentally controlled Okolab cage incubator with stage top chamber set to 37^*°*^ and 85% humidity at low air flow. Time-lapse 3D stacks were acquired with a 10X (0.3 NA) water immersion objective and a Hamamatsu Orca Flash4 camera in single sided stage scan mode, taking a 3D stack every 10 min for 18 h. Image deskewing was performed in SlideBook prior to import into Fiji.

### Image processing in Fiji

Image hyperstacks in native Nikon (.nd2) or Slidebook (.sld) format were imported in Fiji (v2.16.0) ((Schindelin et al. 2012) using BioFormats. Maximum intensity projections of 3D stacks (for each timepoint) were made using the *ZProject* command. To determine hyphal velocity, tips of individual hyphae were identified and tracked using the *Manual Tracking* plugin. For mitochondrial fragmentation, we used *MitochondriaAnalyzer* (Chaudhry et al. 2020).

### Deriving cell numbers from seed density

The number of cells at the start of a time-lapse recording can be accurately determined, as the measured area is directly proportional to the total cell count. Some cells however are close together and show up as clusters ranging from two to ten cells. The exact cell number per cluster was manually determined and added to the calculation. A magnified example is shown in Fig. S2, with A) showing the original micrograph, B) the binarised image after applying adaptive thresholding, and C) the cellular outlines after excluding small, non-cellular fluorescent structures. A total of eight cells can be seen, of which four are aggregated in a cluster. This approach was used for a total of 785 cells, resulting in distributions of cell area and diameter, as shown in Fig. S2D and Fig. S2E. From this analysis, we obtained an estimated mean cell area of 16.8 µm^2^ and a corresponding average diameter of 4.6 µm, consistent with values reported in previous studies (Chavez et al. 2024; Reza et al. 2024). Dividing the total surface area coverage of the basal well by the mean cell area yields a reliable approximation of the initial total number of cells.

### Image processing in MATLAB

We use MATLAB’s image processing toolbox to analyse experimental photographs for the hyphal tracking, area coverage, and biofilm front lightsheet analyses presented in this work. These photographs are in .tiff format. We import photographs into MATLAB as greyscale 16-bit images using imread(). To identify regions occupied by the biofilm, we convert the greyscale images to binary. However, using MATLAB’s `imbinarize()` function directly without prior processing yields poor results. Binarisation underperforms on the original image due to non-uniform illumination in the photograph and lack of contrast between the foreground (biofilm) and the background. To address the issue of non-uniform illumination in the hypha tracking and light sheet datasets, we subtract the background using morphology opening. After obtaining the uniform background, we enhance the contrast of the image using MATLAB’s `imadjust()` function. After adjusting the illumination and contrast, we binarise the image.

When binarising, we use Otsu’s method (`imbinarize()`) for hyphal tracking and biofilm front tracking. This automated thresholding method produces better results than manual threshold selection. For the area coverage, we obtain the most robust results by binarising using an adaptive local mean thresholding approach. Local mean thresholding involves computing the local mean pixel intensity using a mean filter with a 50 *×* 50 pixel window, using `fspecial()` and `imfilter()` in MATLAB. Local mean thresholding is especially important for the corners of images, where the contrast is lower than the centre, even after the pre-processing. We directly apply the local mean threshold method to the greyscale image without pre-processing, as non-uniformity in illumination on the whole-image scale becomes insignificant on the local scale. The local mean thresholding performs better than using pre-processing and Otsu’s method, for which regions occupied by the biofilm near the edge of the photograph are poorly identified. On the other hand, for our photographs pre-processing and Otsu’s method is the best binarisation method for the low-resolution hypha data and light sheet data. For the light sheet data, after binarisation we use Canny edge detection (using the `edge()` MATLAB function) to find the boundary of the largest connected component of the binary image, and this represents the biofilm front.

### Automated Hyphae Tracking Procedure

Hyphal tips are reference points for individual segmented mitochondria fragments. We use an automated image processing method in MATLAB to measure the speed of hyphal growth. This procedure requires us to identify a hyphal tip, and track its position over a time series of photographs. Owing to growth over time, hyphae are easier to identify in later photographs than in earlier photographs. Consequently, we first identify hyphal tips from the final experimental photograph, and measure hyphal speed in reverse order of the experimental photographs, starting from the final photograph. Below is the procedure for tracking the speed of individual hyphae.

1. Binarize all experimental photographs of hyphal growth using the “Image Processing in MATLAB” method described above. For each experiment, we have 15 photographs taken at 10 minute intervals, and we use 60*×* magnification images for hyphal tracking.
2. Identify hyphal tips in the image corresponding to the final time of the experiment. We do this by calculating the maximum Feret diameter for each connected component identified in the binary image, using the `bwferet()` and `bwconncomp()` functions. We then define a hyphal tip to be the point where the maximum Feret diameter meets the edge of the connected component. We denote the position of a hyphal tip (at time *t*_*i*_) as (*x*_*i*_, *y*_*i*_).
3. After identifying hyphal tips, we use the binary image from the next experimental photograph in the time series (proceeding backwards in time) to estimate the hyphal speed. We use the same procedure as Step 2 to identify all hyphal tips on the new image. We then compute the Euclidean distance between the reference tip in the original photograph, and all hyphal tips identified in the new photographs. We assume that the position that corresponds to the same hypha is the tip in the new image that minimises the distance between the tip in the new image and the original tip. We denote this new position as (*x*_*i*_*−* _1_, *y*_*i*_ *−* _1_). The speed, *v*, of the relevant hypha is then approximated using

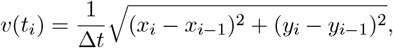

where Δ*t* = 10 minutes is the time interval between two photographs. We then use (*x*_*i*_*−* _1_, *y*_*i*_ *−* _1_) as the new reference position for the next time interval.
4. Repeat Step 3, incrementing backwards one frame at a time, for all hyphal tips identified in the first image in the reversed time series (from the end of the experiment).

We use this procedure to identify and track as many hyphae as possible from a series of experimental photographs. However, to prevent mistracked hyphae, we also apply the following conditions:

1. If the distance between the hyphal tip in any two adjacent frames is greater than 10 µm, then we remove the path from the data set. This is because the time between two frames is approximately 10 minutes, and hyphae are not expected to grow faster than 60 µm h^*−*1^ (Kaneko et al. 2013). Any speeds exceeding this value are likely due to tracking errors.
2. During backtracking, when two hyphae merge, the binarisation procedure will be unable to distinguish the individual hyphae and will instead consider them a single object. If this occurs, then an overlapping path will occur, and two hyphal tips will share the same coordinate. This would cause some tracked paths to be duplicated, so if two paths overlap, one is removed.

For the experiment used in this paper, the automated procedure tracked 181 individual hyphae. This is more hyphae than is typically tracked manually, so the automated procedure yields a larger dataset than manual tracking. The automated tracking takes approximately 1 second per frame, which is also faster than manual tracking.

## Acknowledgements

KL acknowledges funding from the Australian Government through a Research Training Programme Scholarship. SS is supported by a UKRI National Biofilms Innovation Centre Doctoral Training Partnership grant BB/R012415/1. DRP was supported by a National Biofilms Innovation Centre Flexible Talent Mobility Account (BB/S508020/2 IP047). BJB, JEFG, AKYT, and CWG acknowledge funding from the Australian Research Council (Grant numbers DP230100406, DE240100097). PPL and CWG acknowledge equipment funding by the BBSRC (BB/W020033/1).

## Supplementary Material

**Supplementary Fig. S1:**
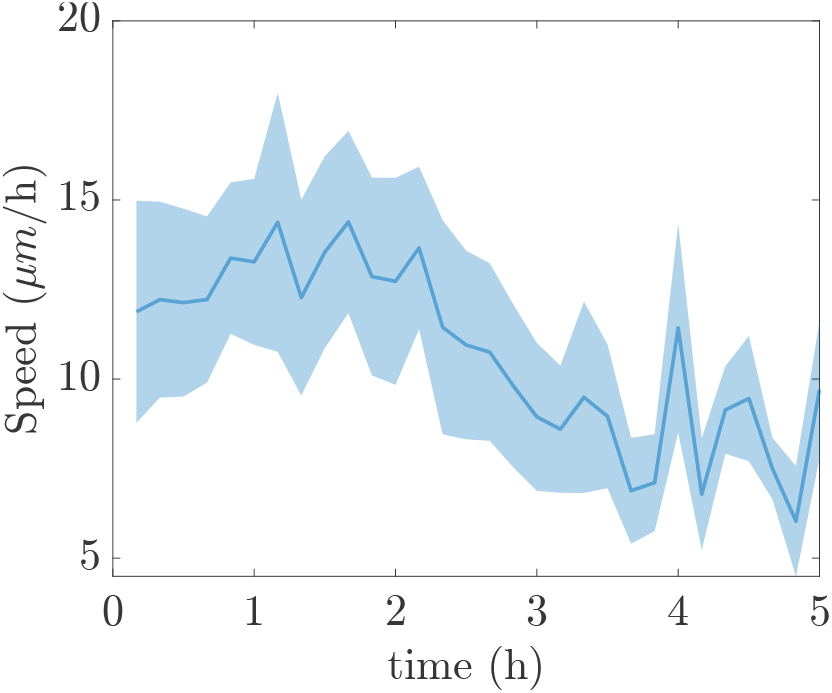
Manually tracked speeds of hyphal emergence for the first five hours, starting with single yeast cells at timepoint 0. The dark blue line represent the mean value of 23 tracks. The light blue area indicates the 95% CI.

**Supplementary Fig. S2:**
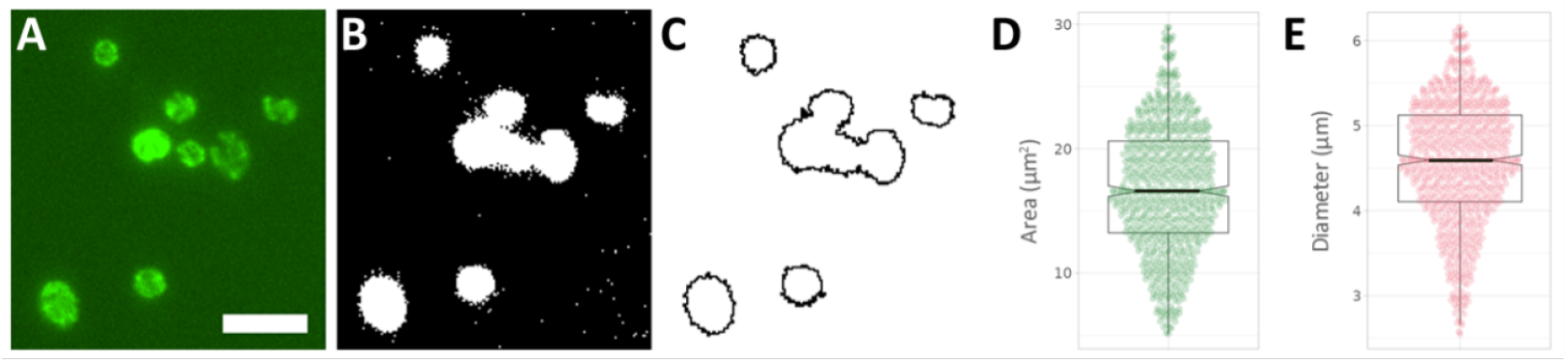
A) Cropped fluorescence micrograph cells of differing sizes. Scalebar 10µm. B) Binarised image after adaptive thresholding. C) Clean outlines for the cells after using size exclusion to remove noise and small, non-cellular particles. D) Boxplot for mean area per cell (16.8 *±* 5.1 µm^2^) with indentations representing the 95% CI. E) Boxplot of corresponding mean cell diameter (4.6 *±* 0.7 µm) with indentations representing the 95% CI.

## References

Cámara, M., W. Green, C. E. MacPhee, P. D. Rakowska, R. Raval, M. C. Richardson, J. Slater-Jefferies, K. Steventon, and J. S. Webb (2022), “Economic significance of biofilms: a multidisciplinary and cross-sectoral challenge”, npj Biofilms and Microbiomes 8, p. 42.

Chaudhry, A., R. Shi, and D. S. Luciani (2020), “A pipeline for multidimensional confocal analysis of mitochondrial morphology, function, and dynamics in pancreatic β-cells”, American Journal of Physiology-Endocrinology and Metabolism 318, E87–E101.

Chavez, C. M., M. Groenewald, A. B. Hulfachor, G. Kpurubu, R. Huerta, C. T. Hittinger, and A. Rokas (2024), “The cell morphological diversity of Saccharomycotina yeasts”, FEMS Yeast Research 24, foad055.

Chen, H., X. Zhou, B. Ren, and L. Cheng (2020), “The regulation of hyphae growth in Candida albicans”, Virulence 11, pp. 337–348.

Duvenage, L., C. A. Munro, and C. W. Gourlay (2019a), “The potential of respiration inhibition as a new approach to combat human fungal pathogens”, Current Genetics 65, pp. 1347–1353.

Duvenage, L., D. R. Pentland, C. A. Munro, and C. W. Gourlay (2019b), “An analysis of respiratory function and mitochondrial morphology in Candida albicans”, BioRxiv, p. 697516.

Gutiérrez–Medina, B. and A. Vázquez-Villa (2021), “Visualizing three-dimensional fungal growth using light sheet fluorescence microscopy”, Fungal Genetics and Biology 150, p. 103549.

Hartmann, R., H. Jeckel, E. Jelli, P. K. Singh, S. Vaidya, M. Bayer, D. K. Rode, L. Vidakovic, F. Díaz-Pascual, J. C. Fong, et al. (2021), “Quantitative image analysis of microbial communities with BiofilmQ”, Nature Microbiology 6, pp. 151–156.

Heydorn, A., A. T. Nielsen, M. Hentzer, C. Sternberg, M. Givskov, B. K. Ersbøll, and S. Molin (2000), “Quantification of biofilm structures by the novel computer program COMSTAT”, Microbiology 146, pp. 2395–2407.

Higuchi-Sanabria, R., T. C. Swayne, I. R. Boldogh, and L. A. Pon (2016), “Live-cell imaging of mitochondria and the actin cytoskeleton in budding yeast”, Cytoskeleton Methods and Protocols: Methods and Protocols, pp. 25–62.

Homann, O. R., J. Dea, S. M. Noble, and A. D. Johnson (2009), “A phenotypic profile of the Candida albicans regulatory network”, PLoS Genetics 5, e1000783.

Hornby, J. M., S. M. Jacobitz-Kizzier, D. J. McNeel, E. C. Jensen, D. S. Treves, and K. W. Nickerson (2004), “Inoculum size effect in dimorphic fungi: extracellular control of yeast-mycelium dimorphism in Ceratocystis ulmi”, Applied and Environmental Microbiology 70, pp. 1356–1359.

Hornby, J. M., E. C. Jensen, A. D. Lisec, J. J. Tasto, B. Jahnke, R. Shoemaker, P. Dussault, and K. W. Nickerson (2001), “Quorum sensing in the dimorphic fungus Candida albicans is mediated by farnesol”, Applied and Environmental Microbiology 67, pp. 2982–2992.

Kaneko, Y., S. Miyagawa, O. Takeda, M. Hakariya, S. Matsumoto, H. Ohno, and Y. Miyazaki (2013), “Real-time microscopic observation of Candida biofilm development and effects due to micafungin and fluconazole”, Antimicrobial Agents and Chemotherapy 57, pp. 2226–2230.

Kiepas, A., E. Voorand, F. Mubaid, P. M. Siegel, and C. M. Brown (2020), “Optimizing live-cell fluorescence imaging conditions to minimize phototoxicity”, Journal of Cell Science 133, jcs242834.

Kumar, A., Y. Wu, R. Christensen, P. Chandris, W. Gandler, E. McCreedy, A. Bokinsky, D. A. Colón-Ramos, Z. Bao, M. McAuliffe, et al. (2014), “Dual-view plane illumination microscopy for rapid and spatially isotropic imaging”, Nature Protocols 9, pp. 2555–2573.

Licea-Rodriguez, J., A. Figueroa-Melendez, K. Falaggis, M. Plata-Sanchez, M. Riquelme, and I. Rocha-Mendoza (2019), “Multicolor fluorescence microscopy using static light sheets and a single-channel detection”, Journal of Biomedical Optics 24, pp. 016501–016501.

Morelli, K. A., J. D. Kerkaert, and R. A. Cramer (2021), “Aspergillus fumigatus biofilms: Toward understanding how growth as a multicellular network increases antifungal resistance and disease progression”, PLoS Pathogens 17, e1009794.

Nagy, G., G. W. Hennig, K. Petrenyi, L. Kovacs, I. Pocsi, V. Dombradi, and G. Banfalvi (2014), “Time-lapse video microscopy and image analysis of adherence and growth patterns of Candida albicans strains”, Applied Microbiology and Biotechnology 98, pp. 5185–5194.

Pentland, D. R., J. Davis, F. A. Mühlschlegel, and C. W. Gourlay (2021), “CO2 enhances the formation, nutrient scavenging and drug resistance properties of C. albicans biofilms”, npj Biofilms and Microbiomes 7, p. 67.

Pentland, D. R., S. Stevens, L. Williams, M. Baker, C. McCall, V. Makarovaite, A. Balfour, F. A. Mühlschlegel, and C. W. Gourlay (2020), “Precision antifungal treatment significantly extends voice prosthesis lifespan in patients following total laryngectomy”, Frontiers in Microbiology 11, p. 975.

Qin, B., C. Fei, A. A. Bridges, A. A. Mashruwala, H. A. Stone, N. S. Wingreen, and B. L. Bassler (2020), “Cell position fates and collective fountain flow in bacterial biofilms revealed by light-sheet microscopy”, Science 369, pp. 71–77.

Ramage, G., S. P. Saville, B. L. Wickes, and J. L. López-Ribot (2002), “Inhibition of Candida albicans biofilm formation by farnesol, a quorum-sensing molecule”, Applied and Environmental Microbiology 68, pp. 5459–5463.

Reza, M. H., S. Dutta, R. Goyal, H. Shah, G. Dey, and K. Sanyal (2024), “Expansion microscopy reveals characteristic ultrastructural features of pathogenic budding yeast species”, Journal of Cell Science 137.

Riquelme, M., J. Aguirre, S. Bartnicki-García, G. H. Braus, M. Feldbrügge, U. Fleig, W. Hansberg, A. Herrera-Estrella, J. Kämper, U. Kück, et al. (2018), “Fungal morphogenesis, from the polarized growth of hyphae to complex reproduction and infection structures”, Microbiology and Molecular Biology Reviews 82, pp. 10–1128.

Schindelin, J., I. Arganda-Carreras, E. Frise, V. Kaynig, M. Longair, T. Pietzsch, S. Preibisch, C. Rueden, S. Saalfeld, B. Schmid, et al. (2012), “Fiji: an open-source platform for biological-image analysis”, Nature Methods 9, pp. 676–682.

Sevilla, M.-J. and F. Odds (1986), “Development of Candida albicans hyphae in different growth media-variations in growth rates, cell dimensions and timing of morphogenetic events”, Microbiology 132, pp. 3083–3088.

Vorregaard, M. (2008), “Comstat2-a modern 3D image analysis environment for biofilms”, MA thesis, Technical University of Denmark, DTU, DK-2800 Kgs. Lyngby, Denmark.

Wong, G. C., J. D. Antani, P. P. Lele, J. Chen, B. Nan, M. J. Kühn, A. Persat, J.-L. Bru, N. M. Høyland-Kroghsbo, A. Siryaporn, et al. (2021), “Roadmap on emerging concepts in the physical biology of bacterial biofilms: from surface sensing to community formation”, Physical Biology 18, p. 051501.

Wu, Y., P. Wawrzusin, J. Senseney, R. S. Fischer, R. Christensen, A. Santella, A. G. York, P. W. Winter, C. M. Waterman, Z. Bao, et al. (2013), “Spatially isotropic four-dimensional imaging with dual-view plane illumination microscopy”, Nature Biotechnology 31, pp. 1032–1038.

